# Insights into the acquisition of the *pks* island and production of colibactin in the *Escherichia coli* population

**DOI:** 10.1101/2021.04.13.439611

**Authors:** Frédéric Auvray, Alexandre Perrat, Yoko Arimizu, Camille V. Chagneau, Nadège Bossuet-Greif, Clémence Massip, Hubert Brugère, Jean-Philippe Nougayrède, Tetsuya Hayashi, Priscilla Branchu, Yoshitoshi Ogura, Eric Oswald

**Affiliations:** IRSD, INSERM, Université de Toulouse, INRA, ENVT, UPS, Toulouse, France; CHU Toulouse, Hôpital Purpan, Service de Bactériologie-Hygiène, Toulouse, France; Department of Bacteriology, Kyushu University, Fukuoka, Japan; Division of Microbiology, Department of Infectious Medicine, Kurume University School of Medicine, Kurume, Fukuoka, Japan

**Keywords:** *pks*, pathogenicity island, genetic diversity, colibactin, genotoxin, *Escherichia coli*, enterobacteria

## Abstract

The *pks* island codes for the enzymes necessary for synthesis of the genotoxin colibactin, which contributes to the virulence of *Escherichia coli* strains and is suspected of promoting colorectal cancer. From a collection of 785 human and bovine *E. coli* isolates, we identified 109 strains carrying a highly conserved *pks* island, mostly from the phylogroup B2, but also from phylogroups A, B1 and D. Different scenarios of *pks* acquisition were deduced from whole genome sequence and phylogenetic analysis. In the main scenario, *pks* was introduced and stabilized into certain sequence types (ST) of the B2 phylogroup, such as ST73 and ST95, at the *asnW* tRNA locus located in the vicinity of the yersiniabactin-encoding High Pathogenicity Island (HPI). In a few B2 strains, *pks* inserted at the *asnU* or *asnV* tRNA *loci* close to the HPI and occasionally was located next to the remnant of an integrative and conjugative element. In a last scenario specific to B1/A strains, *pks* was acquired, independently of the HPI, at a non-tRNA *locus*. All the *pks*-positive strains except 18 produced colibactin. Sixteen strains contained mutations in *clbB* or *clbD*, or a fusion of *clbJ* and *clbK* and were no longer genotoxic but most of them still produced low amount of potentially active metabolites associated with the *pks* island. One strain was fully metabolically inactive without *pks* alteration, but colibactin production was restored by overexpressing the ClbR regulator. In conclusion, the *pks* island is not restricted to human pathogenic B2 strains and is more widely distributed in the *E. coli* population, while preserving its functionality.

**IMPACT STATEMENT:** Colibactin, a genotoxin associated with the carcinogenicity of certain strains of *E. coli*, is encoded by a pathogenicity island called *pks*. We took advantage of a large collection of non-clinical *E. coli* strains originating from human and bovine hosts to explore the distribution, conservation and functionality of the *pks* island. We found that the *pks* island was not only present in the phylogroup B2 (and more specifically to certain B2 sublineages), but also in other genetic phylogroups, highlighting its capacity to disseminate though horizontal gene transfer. We identified various genetic *pks* configurations indicative of an introduction of the *pks* island into *E. coli* on multiple independent occasions. Despite the existence of various acquisition scenarios, we found that the *pks* sequences were highly conserved and *pks*-carrying strains were overwhelmingly capable of producing colibactin, suggesting that the *pks* island is under selective pressure, through the production of colibactin or other secondary metabolites. Future implications include the identification of such metabolites and their biological activities that could be advantageous to *E. coli* and enable its adaptation to various ecological niches.

**DATA SUMMARY:** All sequence data of the 785 *E. coli* used in this study are freely available from the NCBI BioProject database (https://www.ncbi.nlm.nih.gov/bioproject/) under the accession number PRJDB5579. This database was updated to include the sequence data obtained using ONT MinION for the *E. coli* reference strain SP15 and for *E. coli* strains ECSC054, JML285, KS-NP019, NS-NP030 and SI-NP020. The sequence data of *E. coli* strain UPEC129 obtained using PacBio instrument were deposited in the NCBI BioProject database and are available at https://www.ncbi.nlm.nih.gov/Traces/study/ under the accession number PRJNA669570. Hybrid MinION-Illumina and PacBio-Illumina assemblies are available at the NCBI nucleotide database. The genome sequences of 36 other *E. coli* reference strains and 7 non-*E. coli* strains were retrieved from NCBI.

## INTRODUCTION

*Escherichia coli* is not only a commensal resident of the human and animal gut, but also a pathogen responsible for intestinal or extra-intestinal infections. The *E. coli* species is characterized by a high genetic and phenotypic diversity, with a population distributed into at least eight major phylogenetic groups (A, B1, B2, C, D, E, F and G) (1). *E. coli* strains from the phylogroup B2 are increasingly found in the feces of healthy humans in high-income countries and also responsible for extra-intestinal diseases, including urinary tract infections, sepsis, pneumonia and neonatal meningitis (2). By enabling the exchange of genetic material between bacterial cells, horizontal gene transfer (HGT) is a major driving force in the evolution of bacteria, including adaptation to their host and expansion of their ecological niche (3). HGT-mediated acquisition of large genomic islands (GIs) or pathogenicity islands (PAIs) is recognized as a major contributor to the emergence of the various *E. coli* pathotypes (4). The *E. coli pks* pathogenicity island consists of a *clbA*-*clbS* gene cluster enabling the biosynthesis of a polyketide (PK) - non-ribosomal peptide (NRP) hybrid genotoxin known as colibactin (5). This island exhibits typical features of horizontally acquired genomic elements: (i) it is a large (*i*.*e*. 54-kb) region with a distinct GC content compared to that of the chromosomal backbone, (ii) it is physically associated with a phage-type integrase gene that probably mediated its insertion into the chromosome, and (iii) it is located at a tRNA *locus* and is flanked by two short (*i*.*e*. 17-bp) direct repeats (DRs) reminiscent of those generated upon integrase-mediated insertion of mobile genetic elements (5, 6). The *pks* island can be found in other members of *Enterobacteriaceae* such as *Klebsiella pneumoniae, Citrobacter koseri* and *Enterobacter aerogenes* (6), and in the honeybee gut commensal *Frischella perrara* (7) and the marine sponge commensal *Pseudovibrio* sp. (8).

Colibactin is a virulence factor for extra-intestinal pathogenic *E. coli* (ExPEC) (9-11) and is also a suspected procarcinogenic factor (12-14). Colibactin induces DNA interstrand cross-links (ICLs) (15) and double-strand breaks (5) in host eukaryotic cells. Its production involves the sequential action of the Clb proteins, including PK synthases (PKSs), NRP synthetases (NRPSs), hybrid PKS-NRPS and accessory, editing and maturation enzymes (16). Colibactin is first synthetized as a prodrug called precolibactin, carrying an N-myristoyl-D-Asparagine (C14-Asn) side chain that is then cleaved in the periplasm to release the active genotoxin, whose translocation across the bacterial outer membrane remains unknown (17). The production of colibactin is positively regulated by ClbR (18). The multi-modular PKS-NRPS assembly line not only produces colibactin but also a set of numerous secondary metabolites with varying modes of action (19, 20). These include analgesic lipopeptides, such as C12-Asn-GABA, with the capability to diffuse across the epithelial barrier and act on sensory neurons to decrease visceral pain in the host (21). The *pks* island also contributes to the production of siderophores (enterobactin, salmochelin and yersiniabactin), *via* its promiscuous phosphopantetheinyl transferase ClbA (10), and siderophore-microcins *via* its ClbP peptidase (22).

To date, the presence of the *pks* island was investigated mostly in *E. coli* strains isolated from humans with extra-intestinal infections (5, 6, 23, 24). Here we explored the distribution, conservation and functionality of the *pks* island in a large collection of non-clinical *E. coli* strains originating from human and bovine hosts (25). We found that the *pks* island was not only present in the phylogroup B2 but also in other genetic phylogroups. We identified different scenarios for its integration into the *E. coli* genome. The sequence of the *pks* island is highly conserved and *pks*-positive strains were overwhelmingly capable of producing colibactin, suggesting that the *pks* island is under selective pressure for the adaptation of *E. coli* to various ecological niches, through the production of colibactin or other metabolites or *pks*-encoded enzymatic activities.

## METHODS

### Bacterial strains used in the study

The *E. coli* strains were collected in Japan from 418 healthy bovines in 2013 and 2014, 278 healthy humans in 2008, 2009 and 2015, and 89 humans with extra-intestinal infections, either bacteremia (n=67) in 2002-2008 or urinary tract infection (n=22) in 2006 and 2011. They were described recently (25) and corresponded each to a single isolate, duplicates showing less than 5 SNPs difference in their whole genomes being excluded from this study. A list of the 109 *pks*-positive isolates is provided in Table S1. Additional 37 *E. coli* reference strains (Table S2) and 7 non-*E. coli* strains (Table S3) were included in this study; their genome sequences were downloaded from NCBI, except for *E. coli* SP15 which was not available and was obtained here (see below).

### Whole genome sequencing

The whole genome sequences of the 785 *E. coli* isolates were determined by Illumina sequencers (25). Among these, the genomes of 5 *E. coli* strains (ECSC054, JML285, KS-NP019, NS-NP030, and SI-NP020) were further subjected here to long-read sequencing using Oxford Nanopore Technologies (ONT) MinION device. The DNA libraries were prepared using the rapid barcoding kit (Oxford Nanopore Technologies) and sequenced using MinION R9.4.1 flow cells. Long-read sequencing of *E. coli* strain UPEC129 was also performed using Pacific Biosciences (PacBio) RSII sequencer (Genoscreen, Lille, France). The DNA was extracted using Gentra Puregen Yeast/Bact (Qiagen) and the DNA libraries prepared using the SMRTbell Template Prep kit (PacBio). Hybrid assembly of Illumina paired-end reads and MinION or PacBio reads was performed using Unicycler (v.0.4.8) (26). The whole genome sequence of *E. coli* reference strain SP15 was obtained using Illumina and ONT MinION instruments and assembled as described above.

### Sequence and phylogenetic analysis

The core gene–based phylogenetic tree was constructed as described previously (25). Briefly, core genes were determined using Roary (27) and SNP sites were extracted from the core gene alignment using SNP-sites (28). The maximum likelihood (ML) tree was constructed using RAxML (29) with the GTR-GAMMA model and displayed using iTOL (30).

For the phylogenetic analysis of the entire *pks* island, the genome sequences of *pks*-positive strains were aligned with the entire *pks* island sequence of strain IHE3034 using MUMmer (31) and the SNP sites located therein were identified. After removing SNP sites on the VNTR region, a neighbor-joining (NJ) tree was constructed by MEGA7 (32) using the Tamura-Nei evolutionary model.

Cophylogenetic analysis of the core-gene based ML tree and the *pks*-based NJ tree was performed using the “cophylo” function of the R package Phytools (33).

Sequence type and phylogroup determination was performed as described previously (25). The *pks* sequences from four *E. coli* strains belonging to distinct phylogroups (i.e. SI-NP020, KS-NP019, UPEC129 and ECSC054 from phylogroups A, B1, B2 and D, respectively) were extracted from hybrid assemblies and compared at the nucleotide level with that of the reference *E. coli* strain IHE3034. In addition, the amino acid sequences were obtained for the 19 *clb* genes of each strain and aligned by MUSCLE with MEGA7 (32). The alignment file was analyzed with the sequence identity and similarity online software (http://imed.med.ucm.es/Tools/sias.html; accessed in July 2020).

The comparison of *pks* sequences from *E. coli* and other bacterial species was performed with BLASTn (https://blast.ncbi.nlm.nih.gov/Blast.cgi?PAGE_TYPE=BlastSearch). Each *pks* region was defined from *clbA* to *clbS* and used as the query nucleotide sequence against each *pks* region as the subject. Then, the alignment was visualized with Artemis Comparison Tool (v13.0.0) (34).

The integrase nucleotide and amino-acid sequences were aligned using MUSCLE (v3.8.31) and the phylogeny was analysed with PhyML (v3.1/3.0 aLRT) prior to tree visualisation with TreeDyn (v198.3) (http://www.phylogeny.fr; accessed in Sept 2020).

The CC95 strains were typed for their *fimH* allele using FimTyper (v1.0) (https://cge.cbs.dtu.dk/services/FimTyper/; accessed in Nov 2020) and were further assigned to subgroups A-E by analysis of the presence of either of the five subgroup-specific genes described previously (35).

### PCR analysis of the clbJK fusion gene

The 5,651-bp deletion in the *clbJ*-*clbK* region resulting in the *clbJK* fusion gene was tested using a duplex PCR assay, with two primer pairs. The first primer pair (clbK-F, 5’-GACTGCCCAACATACGCTCCG-3’; clbK-R, 5’-TTGTGTCGTTGTACTCTCGGC-3’) was used to amplify a 722 bp-long DNA fragment that is located within the deleted region and is thus only present in strains with an intact *clbJ*-*clbK* region. The second primer pair consisted of primers clbJK-F (5’-AGAATTACCCACTGCCACCA-3’) and clbJK-R (5’-GGCGCTAATGGATCAGATGT-3’) flanking the deleted region, and was used to amplify a 1441 bp-long DNA fragment only present in strains with a *clbJK* fusion gene. The strains with an intact *clbJ*-*clbK* region or a *clbJK* fusion gene yielded a 722-bp or a 1441-bp long amplification product, respectively. Reaction mixture of 50 µL final volume contained 2µL template DNA, 1X GoTaq Reaction buffer, 200µM of each dNTP, 4 mM of MgCl2, 1.25 U of GoTaq DNA polymerase (Promega, France) and 0.2µM of each primer (Eurofins Genomics Ebersberg, Germany). Amplification was done in a GeneAmp® 9700 thermal cycler (Applied Biosystems, Courtaboeuf, France), with the following program: initial denaturation at 95°C for 2 min; 30 cycles of denaturation at 95°C for 30 s, annealing at 56°C for 45 s and extension at 72°C for 1 min 30 s; final extension at 72°C for 5 min. Electrophoresis was carried out in 1% agarose gel and the PCR products visualized after Gel Red (Biotium) staining using a Bio-Rad Chemidoc XRS system (Bio-Rad, France).

### *In vitro* DNA interstrand crosslinking assay

ICL activity was assessed as described previously (15). Briefly, 3 10e6 *E. coli* cells or 6 10e6 *Erwinia oleae* cells pre-grown for 3.5 h in DMEM with 25 mM HEPES (Invitrogen) were mixed with EDTA (1 mM) and 400 ng of linearized plasmid pUC19 DNA and the mixtures were incubated for 40 min at 37°C. After pelleting the bacteria, the DNA was purified from the supernatant and analyzed by electrophoresis on denaturing (40 mM NaOH - 1 mM EDTA) 1% agarose gels. ICL activity of *E. oleae* was also tested in the presence of 400nM 6-histidine-ClbS, which was purified with HisPur nickel-nitrilotriacetic acid (Ni-NTA) agarose (Thermo Scientific) from a culture of BL21(DE3) strain hosting the plasmid pET28a-ClbS-His, as described previously (15).

### Megalocytosis assay

Non-hemolytic *pks*-positive strains were tested for megalocytosis on infected HeLa cells as described previously (5, 36). Briefly, HeLa cells grown to 50% confluence in cell culture 96-well plates were inoculated with 5 µL of overnight culture of bacteria in infection medium (DMEM with 25 mM HEPES) and incubated for 4 h at 37°C in a 5% CO_2_ atmosphere. Cells were then washed and incubated 48 to 72 h in cell culture medium supplemented with 200 µg/mL gentamicin, and then stained with methylene blue for microscopy examination.

### H2AX phosphorylation assay

HeLa cells were infected as described above and H2AX phosphorylation was quantified immediately after the 4 h infection step by immunofluorescence as described elsewhere (37).

### C14-asn quantification

*E. coli* strains were grown for 24 h at 37°C in 10 mL DMEM-HEPES (Gibco), resuspended in 500µL HBSS (Invitrogen) and then crushed with a Precellys instrument (Ozyme, Montigny le Bretonneux, France). After addition of an internal standard mixture (Deuterium-labeled compounds; 400 ng/mL), cold methanol (MeOH) was added and samples were solid-phase extracted on HLB plates (OASIS® HLB 2 mg, 96-well plate, Waters, Ireland). Lipids were eluted with MeOH, evaporated under N_2_, resuspended in MeOH and analysed by high-performance liquid chromatography/tandem mass spectrometry analysis (LC-MS/MS) (MetaToulLipidomics Facility, INSERM UMR1048, Toulouse, France), as described previously (21).

## RESULTS

### The pks island was mainly found in specific E. coli lineages from phylogroup B2

The presence of the *pks* island was investigated in a collection of 785 *E. coli* strains (25) belonging to at least 296 different sequence types (STs) and originating mostly from fecal samples of healthy bovines and humans. Clinical isolates recovered from urine or blood samples of human patients with extra-intestinal infection were also included for comparison. We detected the *pks* island in 109 *E. coli* strains, including 62 (22.3%) out of 278 healthy human fecal isolates and 12 (2.9%) out of 418 healthy bovine fecal isolates (Table 1; Fig. 1). As expected a higher proportion of *pks*-positive strains were found among ExPEC, i.e. 35 (39%) out of 89 strains, including 14 (63.6%) out of 22 strains from urinary tract infection and 21 (31.3%) out of 67 strains from bacteremia. The vast majority of the 109 *pks*-positive strains corresponded to B2 isolates (Table 1; Fig. 1) and the *pks* island was mainly present in specific lineages or STs of the B2 phylogroup (Fig. 1).

**Table 1.** Occurrence of *pks* in *E. coli* strains from healthy humans or bovines, and human patients with extra-intestinal infection.

| Origin | Phylogroup (no. *pks*+ strains / no. strains tested) |  |  |  |  |  |  |  |  |
| --- | --- | --- | --- | --- | --- | --- | --- | --- | --- |
|  | A | B1 | B2 | C | D | E | F | Uncl<sup>c</sup> | Total |
| Healthy bovines<sup>a</sup> | 1/48 | 7/314 | 4/11 | 0/20 | 0/12 | 0/8 | 0/4 | 0/1 | 12/418 |
| Healthy humans<sup>a</sup> | 0/14 | 1/29 | 61/163 | 0/11 | 0/37 | 0/1 | 0/20 | 0/3 | 62/278 |
| Human patients<sup>b</sup> | 0/3 | 0/5 | 34/59 | 0/4 | 1/10 | 0/0 | 0/8 | 0/0 | 35/89 |
| Total | 1/65 | 8/348 | 99/233 | 0/35 | 1/59 | 0/9 | 0/32 | 0/4 | 109/785 |

<sup>a</sup> Isolates were collected from feces of healthy individuals <sup>b</sup> Isolates were collected from blood (n=67) or urine (n=22) from human patients with extra-intestinal infection. <sup>c</sup> Unclassified

**Figure 1.**
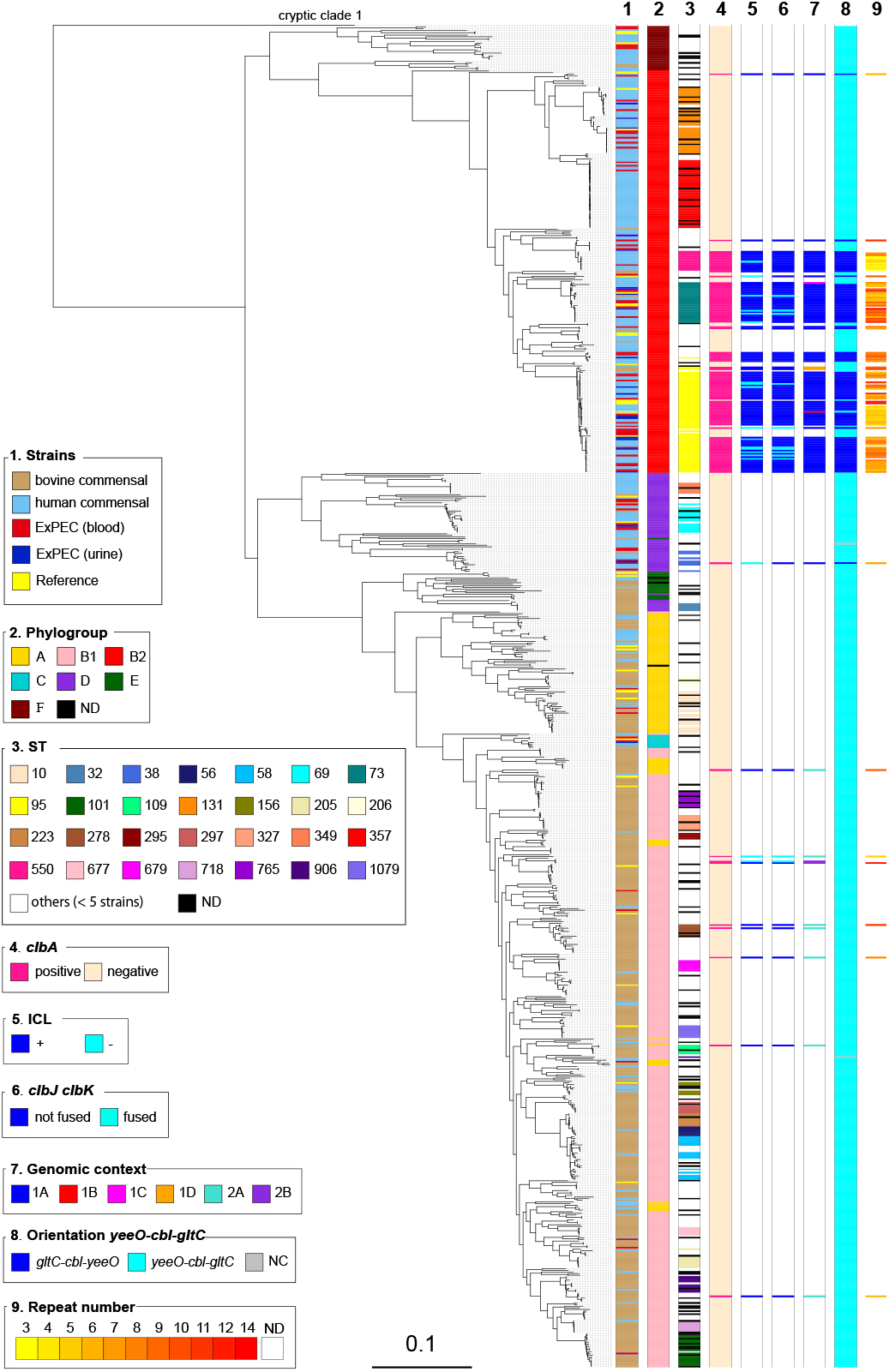
Phylogenetic relationship and distribution of *pks*-positive/negative *E. coli* isolates among 696 human and bovine commensal *E. coli*, 89 ExPEC and 37 completely sequenced reference *E. coli* strains. A core gene-based maximum likelihood (ML) tree was constructed based on 271,403 SNPs located on 2,000 core genes and rooted on cryptic *Escherichia* clade 1 strains as outgroups. Origin (column 1), phylogroup (column 2), major sequence type (ST) (*i*.*e*. ST identified for at least 5 strains) (column 3), presence of *pks* (*clbA*) (column 4), colibactin activity (ICL) (column 5), presence of the *clbJK* fusion gene (column 6), genetic *pks* configuration (see Fig. 3) (column 7), orientation of the *asnV-asnU-asnW* region situated downstream the *asnT* tRNA gene (column 8) and the number of repeats 5’-ACAGATAC-3’ found in the *clbB*-*clbR* intergenic region (see Fig. 2) (column 9) are shown for each strain. ND, not determined.

Strikingly, the *pks* island was found in (nearly) 100% of strains belonging to ST12, ST73, ST95 and ST550, while it was excluded from other STs, such as ST131 and ST357 (Fig. 1; Table 2). Interestingly, these *pks*-positive and -negative STs are found in distinct clusters in the core-genome based phylogenetic tree (Fig. 1) suggesting that *pks* acquisition occurred after the divergence of these clusters from a common ancestor. We further characterized the 54 *pks*-positive strains of ST95 for their *fimH* allele and affiliation to CC95 subgroups A to E defined previously (35). We could assign 35 of them to subgroup A (n=22), B (n=12) or E (n=1) (Table S1). The remaining 19 strains, including 15 of serotype O1:H1, did not belong to any of these five subgroups. No *pks*-positive strain was assigned to CC95 subgroups C or D, in agreement with previous results (35).

**Table 2.**
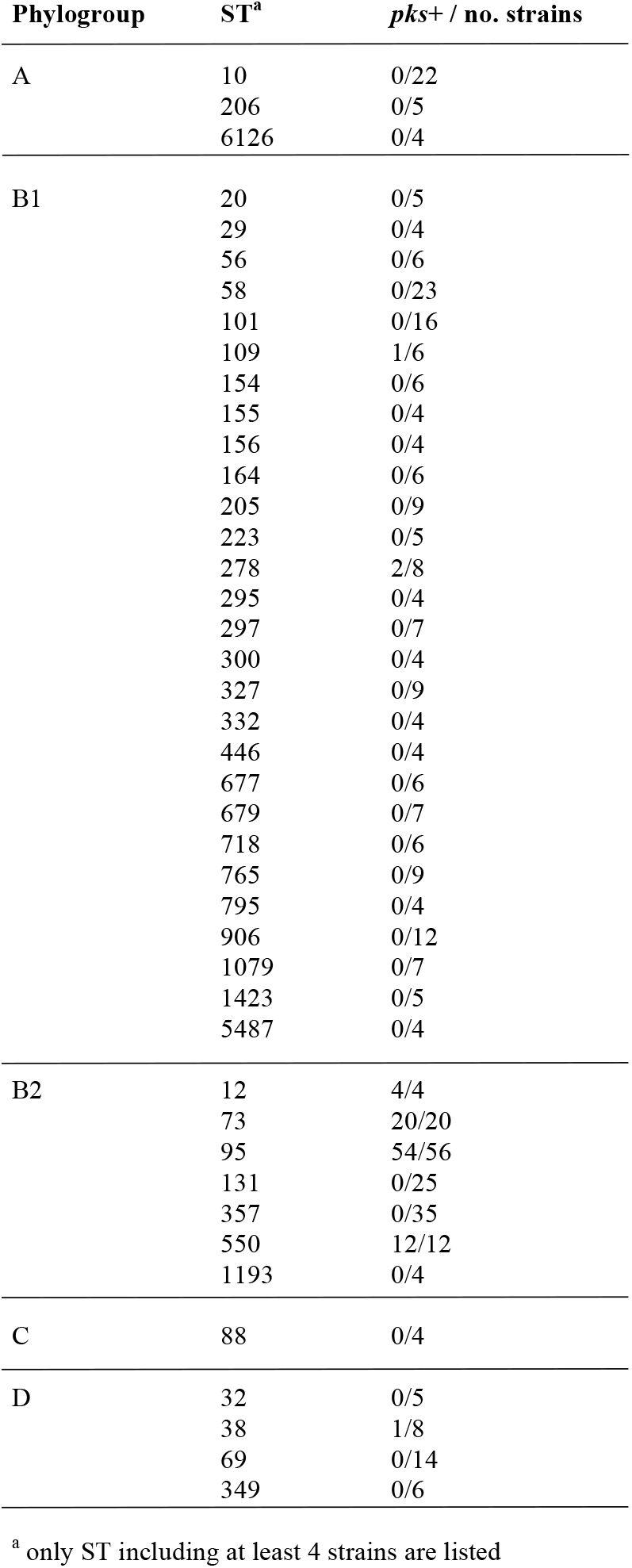
Distribution of the *pks* island in the predominant sequence types (ST) among *E. coli* strains isolated from healthy bovines, healthy humans and human patients with extra-intestinal infection.

Except for four B2 strains originating from healthy bovines, the *pks*-positive B2 isolates originated from humans, either patients with extra-intestinal infection (n=34) or healthy individuals (n=61) (Fig. 1; Table 1). The low occurrence of *pks* among bovine isolates likely reflected the low prevalence of B2 strains in cattle (25). Interestingly, 10 non-B2 *pks*-positive strains were identified corresponding to 1 human blood isolate from phylogroup D, 1 healthy human fecal isolate from group B1 and 8 healthy bovine fecal isolates from groups A (n=1) and B1 (n=7) (Fig. 1, Table 1). In contrast to the B2 *pks*-positive isolates, these strains were scattered throughout the core genome phylogenetic tree and were not representative of any particular lineage or ST (Fig. 1; Table 2).

### High level of genetic conservation of the pks island among E. coli phylogroups and other enterobacteria

The *pks* sequence from the B2 reference *E. coli* strain IHE3034 was compared to that of three non-B2 *E. coli* isolates, including the single group A isolate (i.e. SI-NP020), the single group D isolate (ECSC054) and one out of the eight B1 isolates (i.e. KS-NP019). An additional B2 isolate (i.e. UPEC129) was also selected for this analysis. To perform this comparison, the whole genomes of these four isolates were assembled from a combination of short and long reads. At the amino acid level, over 99 % identity was observed for each of the 19 *clb* gene products (Fig. 2). At the nucleotide level, the only variation observed in the *pks* sequence was the size of the region located between *clbB* and *clbR* which contains a variable number of tandem repeats (VNTR) of the motif 5’-ACAGATAC-3’ (6). This VNTR locus contained between 3 and 14 repeat units when the whole collection of *pks*-positive strains was analysed (except for 17 isolates for which the VNTR length could not be calculated), with no apparent correlation with the STs (Fig. 2). Therefore, apart from the size of the VNTR, the *pks* island was highly conserved among the strains, irrespective of their phylogroup or ST.

**Figure 2.**
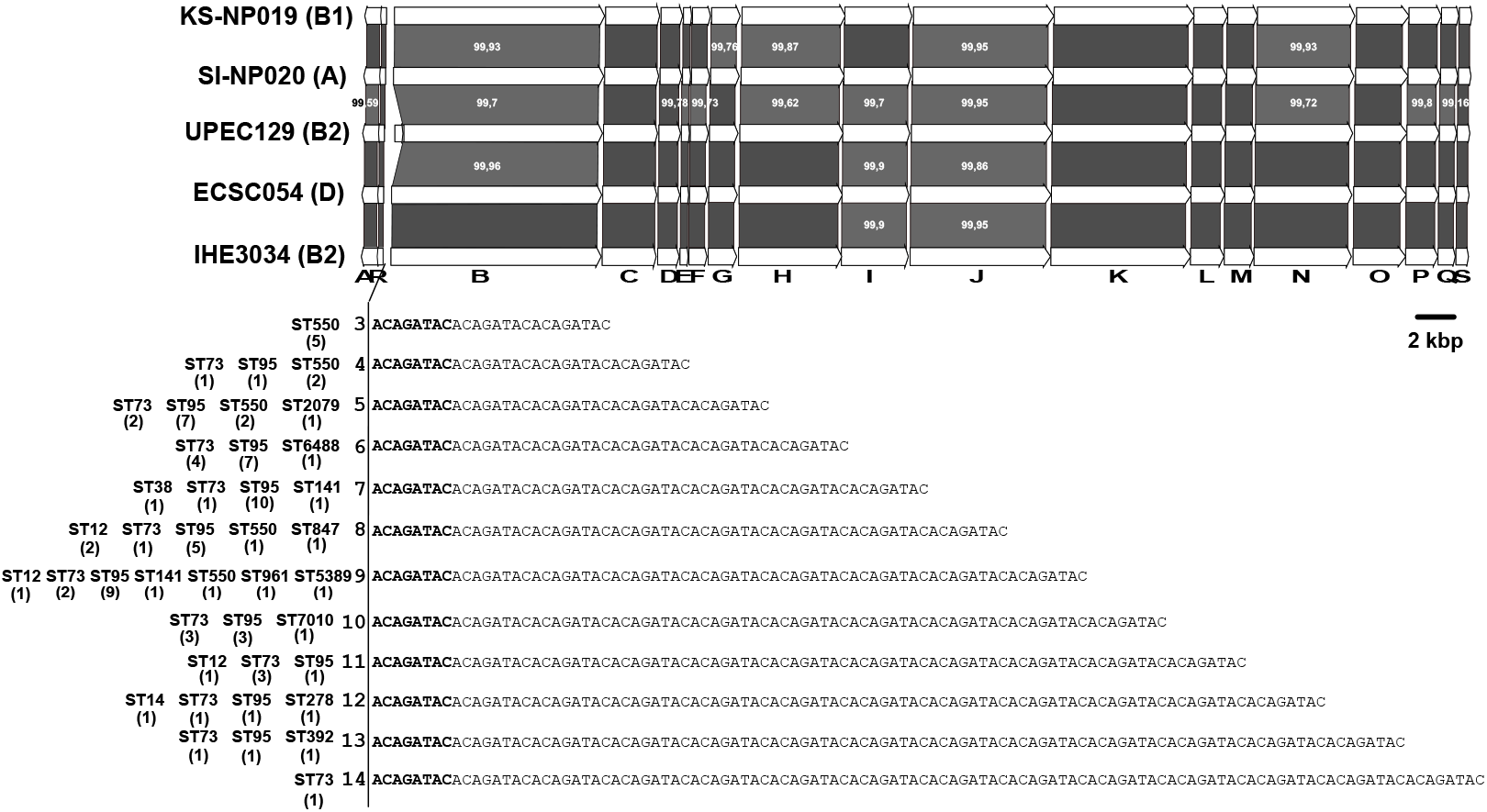
Comparison of the *pks* islands of *E. coli* strains belonging to phylogroups A, B1, B2 and D. The 19 ORFs of the *clbA*-*clbS* gene cluster from the reference *E. coli* strain IHE3034 sequence (group B2) and MinION- or PacBio-derived sequences of *E. coli* strains KS-NP019 (group B1), SI-NP020 (group A), UPEC129 (group B2) and ECSC054 (group D) are represented by arrows with the arrowhead representing direction of transcription. The areas between the corresponding genetic maps shaded in dark and light gray indicate 100% amino acid identity and *ca*. 99% amino acid similarity, respectively. The number of repeated motif 5’-ACAGATAC-3’ found in the *clbB*-*clbR* nucleotide intergenic region of *pks*-positive *E. coli* strains and the sequence types (ST) of the corresponding strains are indicated below the *clbA*-*clbS* gene cluster, with the number of strains into parenthesis.

Comparison of the *pks* island nucleotide sequence from B2 reference strain IHE3034 with that of other *pks*-positive bacterial species confirmed that it was conserved in other members of the *Enterobacteriaceae* (Fig. S1) such as *K. pneumoniae, E. aerogenes, C. koseri, Serratia marcescens* and *Erwinia oleae*. A similar *pks* island was present, although less conserved in *F. perrara* and *Pseudovibrio* sp. (Fig. S1).

### The pks islands in E. coli from phylogenetic groups B2 and D share a similar genomic environment

To gain insights into the events leading to the acquisition of the *pks* island into the *E. coli* population, we analysed the genomic environment of the *pks* island in the 109 *pks*-positive strains and in seven *pks*-positive *E. coli* reference strains (536, ABU83972, CFT073, Nissle 1917, UTI89, IHE3034 and SP15). Various configurations were found for the *pks* island genomic environment, suggesting two main scenarios of *pks* acquisition, depending on the presence or absence of an integrase gene (Fig. 3). The genetic configuration typical of B2 strains, named 1A, which is characterized by a *pks* island carrying an integrase gene and inserted into the *asnW* tRNA gene in the vicinity of the *asnT*-located High Pathogenicity Island (HPI) (5, 6) was found in 96 B2 strains of our collection and in one phylogroup D strain, ECSC054 (Fig 1; Fig. 3).

**Figure 3.**
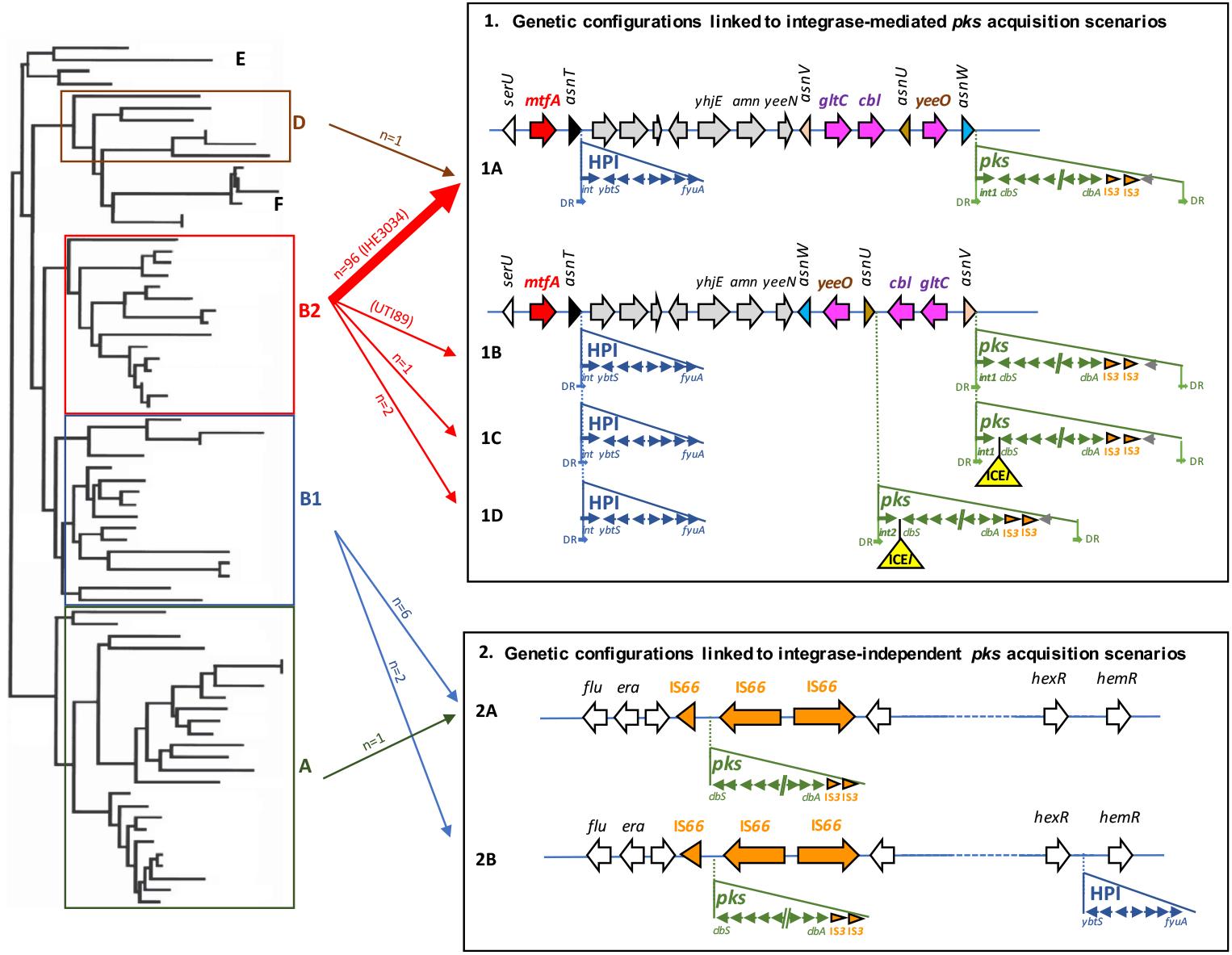
Genetic configurations of the *pks* island and HPI in *E. coli* strains and proposed scenarios for their acquisition. **Left**. Schematic phylogenetic tree showing the distribution of *E. coli* into the main phylogroups. **Right**. *E. coli* genetic *pks* and HPI configurations resulting from proposed acquisition scenarios involving site-specific recombination (configurations 1A, 1B, 1C and 1D) or not (configurations 2A and 2B). The location and orientation of the tRNA genes and the ORFs of the chromosomal regions, including the integrase and genes from *pks* and the HPI, are indicated by the arrows. Partial and complete IS elements are represented by orange arrowheads and arrows, respectively. The ICE-*like* element (ICE*l*) found in configurations 1C and 1D is represented as a yellow triangle. DR, direct repeats located at the extremities of the islands (except for the HPI in configurations 1A-1D, one DR lacking at the right border). **Middle**. The arrows connect the phylogroups A, B1, B2 and D (left) with the *pks* and HPI configurations (right). The number of *E. coli* isolates belonging to the collection of 785 strains and corresponding to each configuration are indicated (except for configuration 1B which was only found in reference strain UTI89 indicated in parenthesis). The thick arrow represents the most frequently found configuration (exemplified here by reference strain IHE3034 indicated in parenthesis).

A similar configuration was found in a few other B2 strains but with variations in the location of the *pks* island which was inserted either into the *asnV* (corresponding to configurations 1B and 1C found in ST95 reference strain UTI89 and in ST73 strain JML226, respectively) or *asnU* tRNA gene (corresponding to configuration 1D found in strains KS-P003 and KS-P027, both belonging to ST95) (Fig. 1; Fig. 3). Besides the difference in the tRNA insertion site, the configuration 1B found in reference strain UTI89 differed from the major configuration 1A by the orientation of the 4,309-bp *asnW-asnU-asnV* tRNA region upstream of the *pks* island. This region contains three other genes, namely *gltC* and *cbl*, encoding two LysR-family transcriptional regulators, and *yeeO*, encoding a flavin mononucleotide [FMN] and flavin adenine dinucleotide [FAD] exporter. The configurations 1C and 1D possessed the same *asnW-asnU-asnV* orientation as in UTI89 and carried a 25-kb region between the *pks* integrase gene and *clbS*. A 14-kb section from this region exhibited high sequence similarity (>99%) to integrative and conjugative elements (ICE) identified in *E. coli* (ICE*Ec1*) and *K. pneumoniae* (ICE*Kp1*), in particular to the DNA regions I and II from ICE*Ec1* involved in mating-pair formation (Mpf) and DNA mobilization, respectively (Fig. 4) (6, 38). This 14-kb section could therefore be considered as an ICE-like element, although it is most likely non-functional given the lack of a complete region II (Fig. 4). The remaining 11-kb section was not homologous to ICE*Ec1*, ICE*Kp1* or any other ICE, and its role could not be predicted.

**Figure 4.**
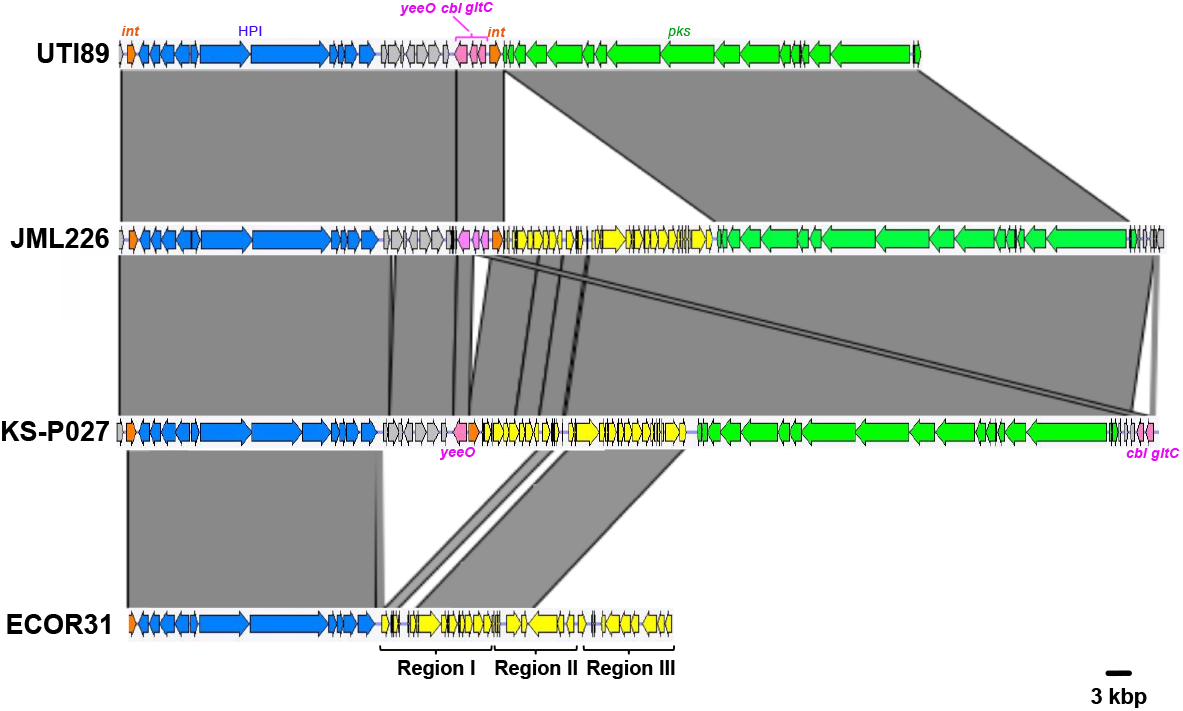
Comparison of the chromosomal region covering the HPI and *pks* island between 3 atypical *E. coli* B2 strains (UTI89, JML226 and KS-P027, with genetic configurations 1B, 1C and 1D, respectively) and the integrative conjugative element ICE*Ec1* from *E. coli* strain ECOR31. Nucleotide sequence similarity (>99%) between different DNA regions is indicated by grey areas between the corresponding genetic maps. The *pks* island and the HPI are represented in green and blue, respectively, and the integrase genes in orange. The *yeeO, cbl* and *gltC* genes located between the *asnV* and *asnW* tRNA genes in UTI89 are represented in pink. The region between the HPI and the *yeeO* gene is represented in grey and the ICE-related region inserted either next to *pks* (JML226 and KS-P027) or next to the HPI (ECOR31) is represented in yellow. In the ECOR31 strain, the ICE is divided in three parts, including region I encoding a mating pair formation system, region II encoding a DNA-processing system, both involved in conjugative transfer, and region III comprising hypothetical genes.

The phage-type *pks* integrase is a tyrosine site-specific recombinase with similarity to the phage P4 integrase C-terminal catalytic domain (INT_P4_C). The integrase genes located at the *asnW* (configuration 1A) or *asnV* loci (configurations 1B and 1C) and their gene products were highly conserved and grouped into the integrase family 1 (Fig. S2), whereas the integrase genes located at the *asnU* locus (configuration 1D) and their gene products shared 94% nucleotide and 94.6% amino acid sequence similarity, respectively, with those of family 1 and were thus grouped into the integrase family 2 (Fig. S2).

### Atypical genomic environments of pks islands in E. coli from phylogenetic groups A and B1

In the 9 *pks*-positive *E. coli* strains from phylogroups A and B1, two different configurations (named 2A and 2B) were observed that drastically differed from those found in B2/D strains. Their *pks* islands lacked an integrase gene, were not inserted into a tRNA gene and there were no direct repeats at their chromosomal boundaries (Fig. 3). The *pks* islands were located in the vicinity of the genes *flu* (or *agn43*) and *era* encoding the Ag43 autotransporter adhesin and a GTPase essential for cell growth and viability, respectively. They were flanked on one side by a truncated copy of the IS*66* insertion sequence (IS) and on the other side by two intact IS*66* copies. Two truncated copies of IS*3* were also found next to the *clbA* gene but this was also the case for configurations 1A-1D. Moreover, in these B1/A strains, the HPI was absent (Table 3; Fig. 3, configuration 2A), except for two isolates in which the HPI was present but not in the vicinity of the *pks* island and not into a tRNA *locus* (Table 3; Fig. 3, configuration 2B). Using PCR assays, it was shown previously that three *E. coli* strains from phylogroup B1 (namely U12633, U15156 and U19010) possessed a *pks* island that co-localized with the HPI and the DNA transfer and mobilization region of an ICE*Ec1*-like element (6), a situation that is reminiscent of that of configuration 1D. However, as the whole genome sequences of these three strains were not available, this could not be confirmed here.

**Table 3.**
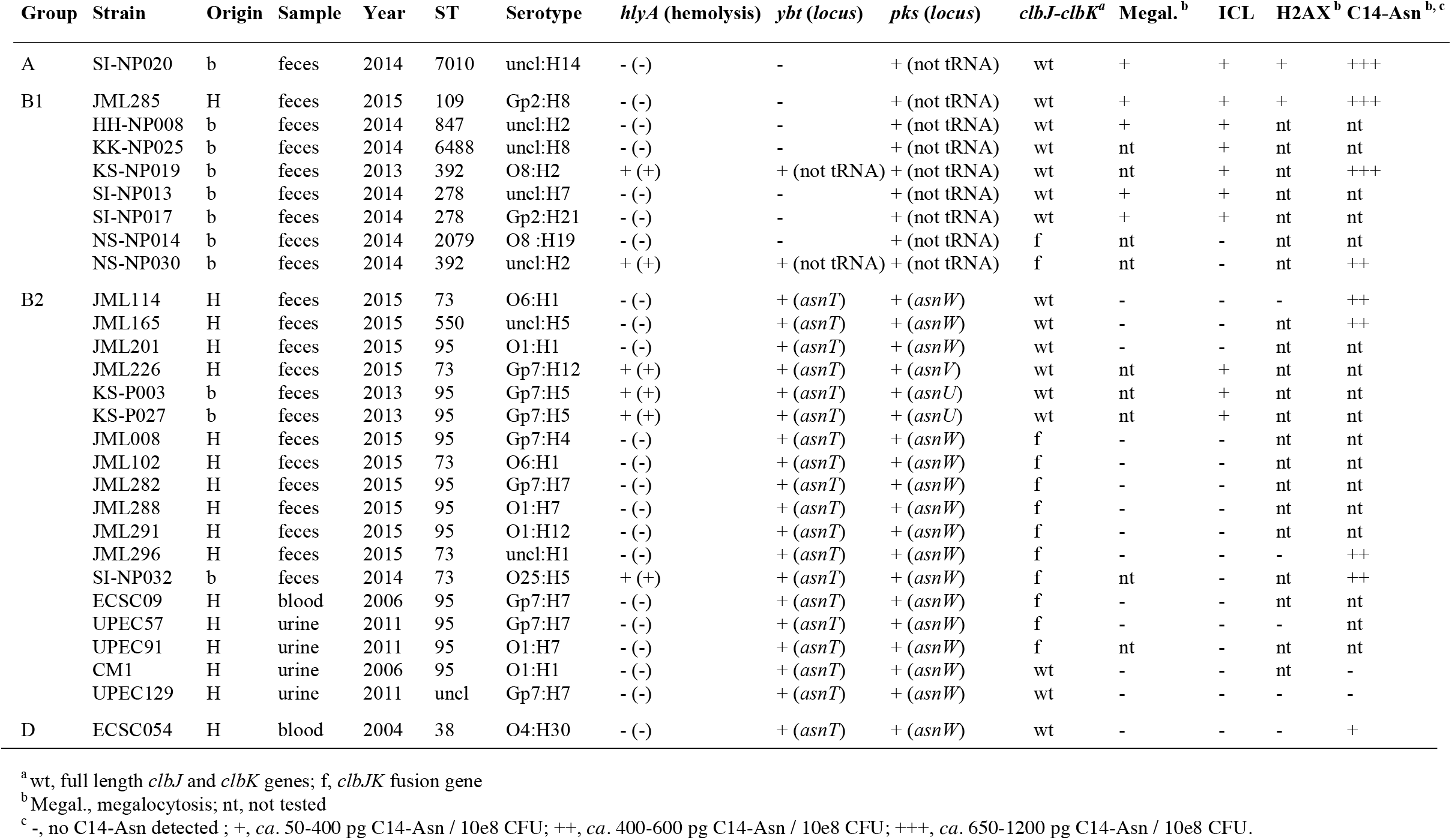
Characteristics of B2 and non-B2 *pks*-positive *E. coli* strains with atypical features regarding *pks* integrity, functionality or location.

Since the *asnW-asnU-asnV* tRNA region displayed distinct orientations in *pks*-positive B2 strains depending on *pks* configuration, we further analysed its orientation for the rest of the *E. coli* collection, i.e. in *pks*-positive B1/A strains and in *pks*-negative strains. The “*asnV-asnU-asnW”* orientation was uniquely found in typical *pks*-positive B2 strains with configuration 1A, suggesting that, in these strains, *pks* acquisition at the *asnW* locus was accompanied by an inversion of the upstream tRNA-encoding region (Fig. 1; Fig. 3).

### The phylogeny of the pks island globally reflects that of the E. coli core genome

To shed further lights on the different *pks* acquisition scenarios, we constructed a phylogenetic tree of the entire *pks* sequences (i.e. from *clbA* to *clbS*, except for the VNTR-containing region which was excluded from the analysis) from the 109 *pks*-positive strains. Globally, the *pks* sequences from the strains showing distinct *pks* genomic configurations formed distinct clusters (Fig. 5). The *pks* sequence of strain UTI89 with a unique configuration (configuration 1B) was clustered with those of strains with configuration 1A (Fig. 5). Remarkably, the *pks* sequences of B1/A strains segregated separately from those of B2/D strains. Moreover, the *pks* sequences with an insertion of an ICE-like element in the B2 (ST73) human strain JML226 (with configuration 1C) or in the pair of B2 (ST95) bovine strains KS-P003 and KS-P027 (with configuration 1D) also clustered separately and were closer to the *pks* sequences of B1/A strains than to those of the B2/D strains lacking this ICE-like element (Fig. 5). Finally, the *pks* sequences of *C. koseri, E. aerogenes, K. pneumoniae* and *S. marcescens* were close to those of *E. coli* B2/D strains with configuration 1A (Fig. 5) whereas that of *E. oleae* was more phylogenetically distant and clustered separately (data not shown).

**Figure 5.**
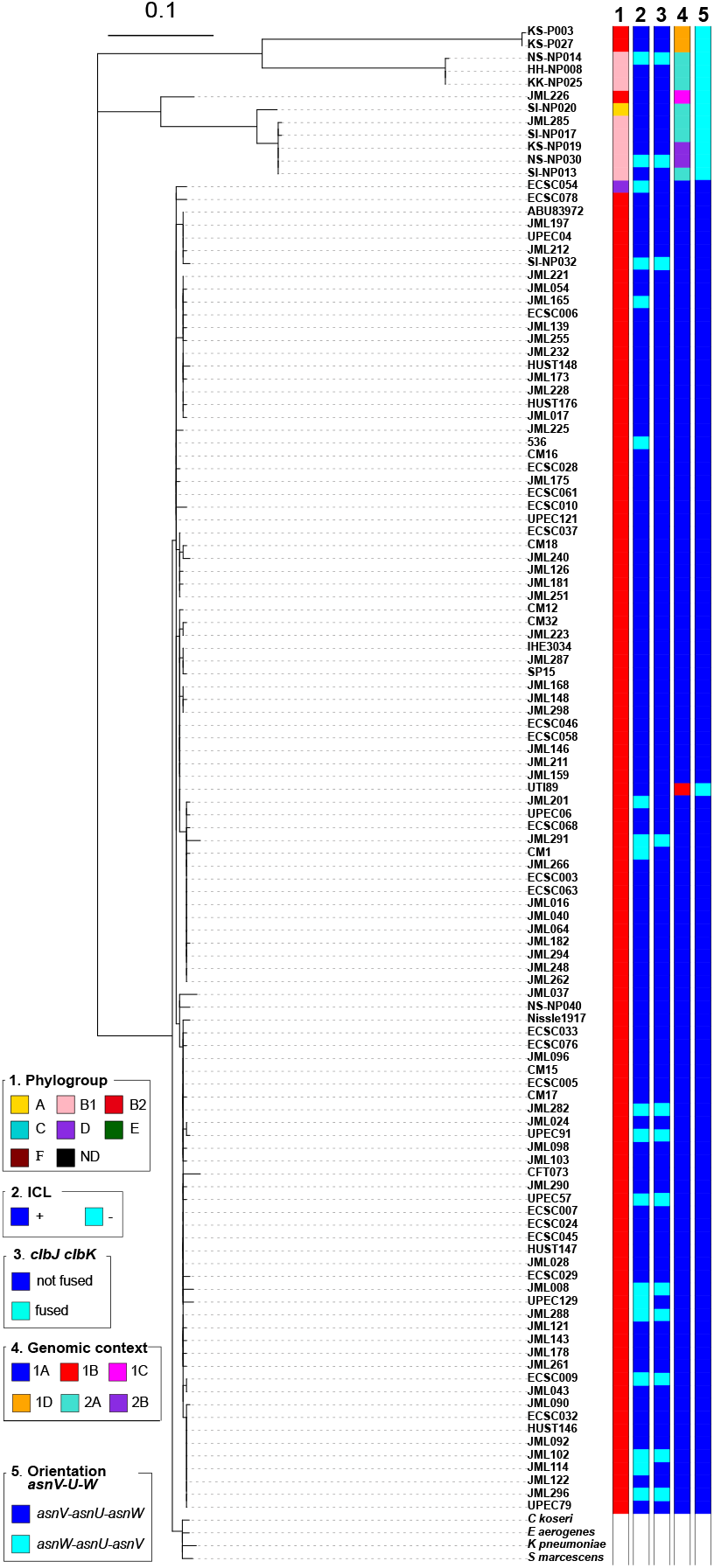
Phylogenetic tree of the entire *pks* island. SNP analysis was performed with the *pks* sequences of the 109 *pks*-positive *E. coli* strains and a NJ phylogenetic tree was built. The *pks* sequences from 7 reference *E. coli* strains (536, ABU83972, CFT073, Nissle 1917, UTI89, IHE3034 and SP15) and other enterobacteria (*i*.*e. C. koseri* ATCC BAA-895, *E. aerogenes* EA1509E, *K. pneumoniae* 1084, and *S. marcescens* AS012490) were also included in this tree. Phylogroup (column 1), colibactin activity (column 2), presence of a *clbJK* fusion gene (column 3), genetic *pks* configuration (see Fig. 3) (column 4) and orientation of the *asnV-asnU-asnW* region (see Fig. 3) (column 5) are indicated for each *pks*-positive strain.

To further assess the evolutionary relationships between the *pks* sequences and the genetic background of the strains, a cophylogenetic analysis was performed where the phylogenetic trees based on the *pks* sequence and the core genome were compared. Globally, congruence was observed between both trees (Fig. 6). It was noticeable that most of the typical *pks*-positive B2 strains whose core genomes clustered together into lineages of clonal complexes (CC) 12, CC14, CC73 and CC95 contained *pks* sequences that also clustered together in different subgroups of the main *pks* cluster (Fig. 6). In particular, the CC95 strains that clustered together into subgroups A and B (as defined by *fimH* typing) or O1:H1 subgroup in the core genome tree also clustered together in the *pks* tree (Fig. 6). These observations support the hypothesis of an introduction of the *pks* island into CC12, CC14, CC73 and CC95 through horizontal acquisition by their most recent common ancestor (MRCA) or by the MRCA of each of these lineages, followed by vertical transmission with subtle *pks* divergence overtime. Since CC95 subgroups C or D contain *pks*-negative strains (35) (strain ECSC026, subgroup C; this study), we further hypothesize that *pks* was lost during the evolution of these sublineages.

**Figure 6.**
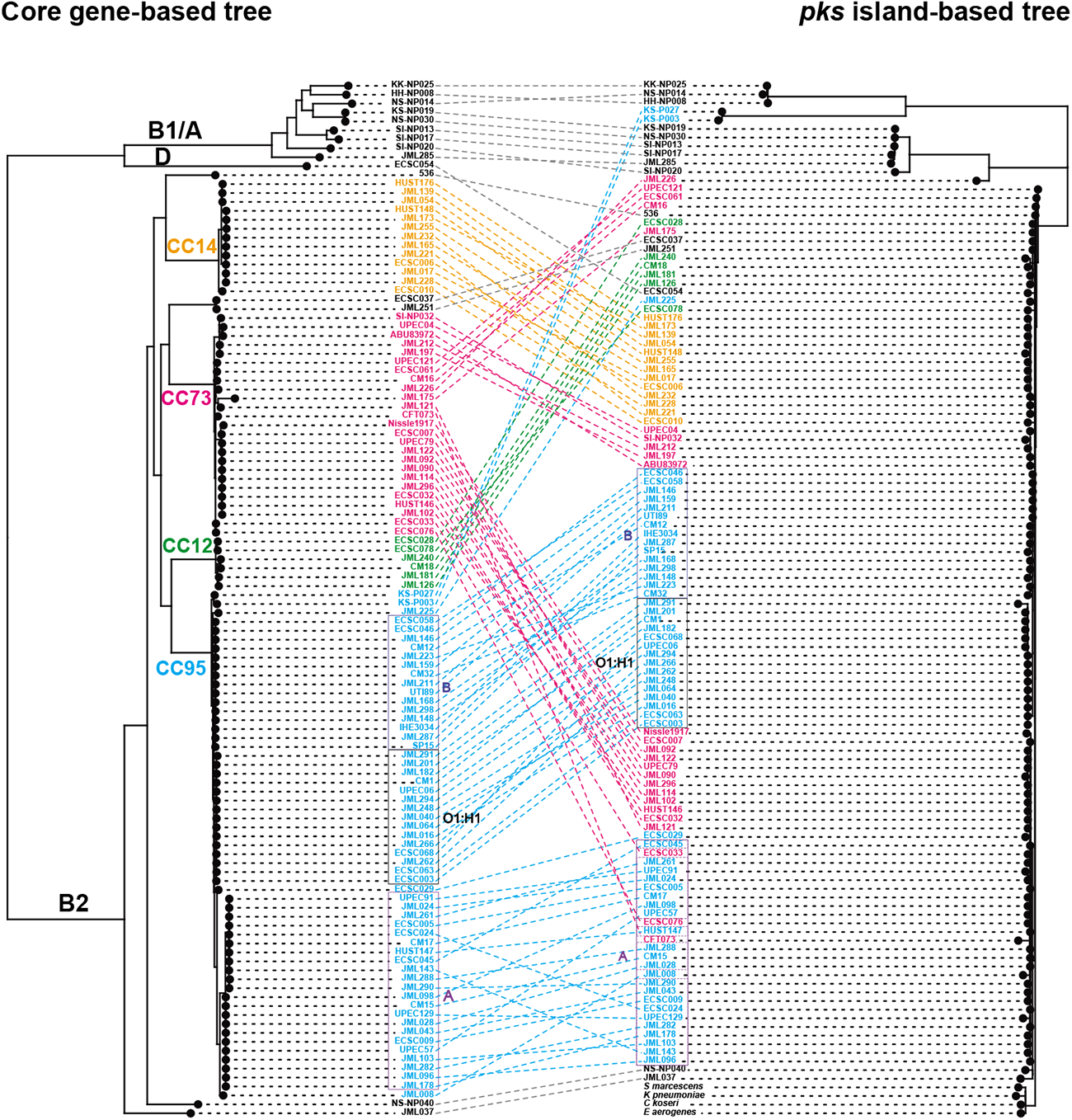
Cophylogeny of *pks* sequences and *E. coli* host strains. A comparison generated with Phytools of *E. coli* core gene-based ML tree and *pks*-based NJ tree is shown, including the links between *pks* and host strains (dashed lines). The phylogenetic groups (A, B1, B2 and D) are indicated. The strains belonging to the major clonal complexes (CCs) are shown with coloured names, including those of CC12 (containing 4 ST12 strains, one ST961 strain [ECSC078] and one ST5389 strain [ECSC078]), CC14 (containing one ST14 strain [ECSC010] and 12 ST550 strains), CC73 (containing only ST73 strains) and CC95 (containing only ST95 strains). Strains from CC95 that belong to subgroups A and B (as defined by *fimH* typing) or to serotype O1:H1 are boxed. 7 reference *pks*-positive *E. coli* strains (536, ABU83972, CFT073, Nissle 1917, UTI89, IHE3034 and SP15) were included in both trees, whereas the non-*E. coli* strains carrying a *pks* island (i.e. *C. koseri* ATCC BAA-895, *E. aerogenes* EA1509E, *K. pneumoniae* 1084, and *S. marcescens* AS012490) were included only in the *pks*-based tree.

The fact that a single *pks*-positive strain from phylogroup D (ECSC054) possessed a *pks* island whose sequence clustered with that of B2 strains (Fig. 6) suggests that this strain acquired *pks* from a B2 strain through HGT. The *pks*-carrying B1/A strains were diverse based on their core genomes and their *pks* sequences clustered in two separate groups that were distantly related to the major *pks* cluster of B2 strains (Fig. 6), suggesting the existence of sporadic *pks* introduction within the B1 and A phylogroups, presumably through HGT from a donor strain different from typical *pks*-positive B2 strains. Finally, the cophylogeny also confirmed that the two atypical B2 ST95 strains KS-P03 and KS-P027 clustered with the other B2 strains of ST95 based on the core genome but contained a divergent *pks* sequence which was closer to those of B1 or A strains (Fig. 6), suggesting that this pair of strains probably acquired their *pks* islands through HGT, possibly from a donor strain carrying a *pks* island with an ICE insertion. The same scenario also presumably occurred with the atypical B2 ST73 strain JML226 which clustered with the other B2 strains of ST73 in the core genome tree but carried a *pks* island characterized by an ICE-like insertion and a sequence closer to those of B1/A strains than to those of B2 ST73 strains.

### The functionality of the cluster of genes of the pks island is conserved in the majority of the enterobacterial strains

We next investigated the functionality of the *pks* islands in *E. coli* strains belonging to various phylogroups and carrying phylogenetically distinct *pks* sequences, as well as in the *E. oleae* strain DAPP-PG531. The production of the genotoxin colibactin was directly investigated through the formation of DNA interstrand cross-links (ICLs) (Fig. S3A). The vast majority of the *E. coli* strains carrying the *pks* island (i.e. 83.5%) produced ICLs (Fig. 1). DNA-crosslinking was also observed for the *E. oleae* strain, and it was abrogated by adding purified colibactin self-resistance protein ClbS (Fig. S4), confirming the production of a *bona fide* colibactin by this strain carrying a less conserved sequence of the *pks* island. The *E. coli* genotoxic strains belonged to phylogroups B2, B1 and A (Fig. 1). Eighteen (16.5%) *pks*-positive *E. coli* isolates lacked a detectable interstrand crosslinking activity, including 15 strains from phylogroup B2, 2 strains from phylogroup B1 and the single *pks*-positive strain from phylogroup D (Table 3). These strains did not cluster together in the core genome phylogenetic tree but instead were intertwined among genotoxic strains (Fig. 1, Fig. 5). To confirm the absence of genotoxicity, we tested the ability of these ICL-negative strains to trigger megalocytosis in cultured HeLa cells (Fig. S3B) and phosphorylation of histone H2AX (Fig. S3C), a robust marker for DNA damage in eukaryotic cells. To avoid cell lysis during infection, we assessed only non-hemolytic strains. No megalocytosis and no p-H2AX foci were detected in HeLa cells exposed to subsets of ICL-negative strains (Table 3; n=14 and n=5, respectively), even at a high multiplicity of infection, confirming the deficiency of these strains in colibactin production. These results showed that except for a few strains, *E. coli* strains carrying a *pks* island are overwhelmingly capable of producing the genotoxin colibactin, regardless of their phylogenies and genomic configurations of their *pks* islands.

To examine the reasons for the lack of genotoxic activity of the 18 ICL-negative *E. coli* strains, we further analyzed the sequence of the *pks* island from those strains. We identified genetic alterations of the *pks* island in 16 out of the 18 non-genotoxic isolates. Strain JML114 carried a single nucleotide deletion in *clbD* at position 172 (A), leading to the segregation of *clbD* into two ORFs (Fig. 7). Strain JML165 carried an IS*1* inserted at the 3’-end of *clbD* after position 838. In strains UPEC129 and JML201, *clbB* was segregated into two ORFs due to nucleotide substitutions at positions 452 and 453 (AC to GA) (UPEC129; Fig. 7), and at position 872 (G to A) (JML201; data not shown), respectively. The genetic alterations identified in these four non-genotoxic strains each resulted in premature stop codons in *clbB* or *clbD* genes coding enzymes that are essential for the production of colibactin.

**Figure 7.**
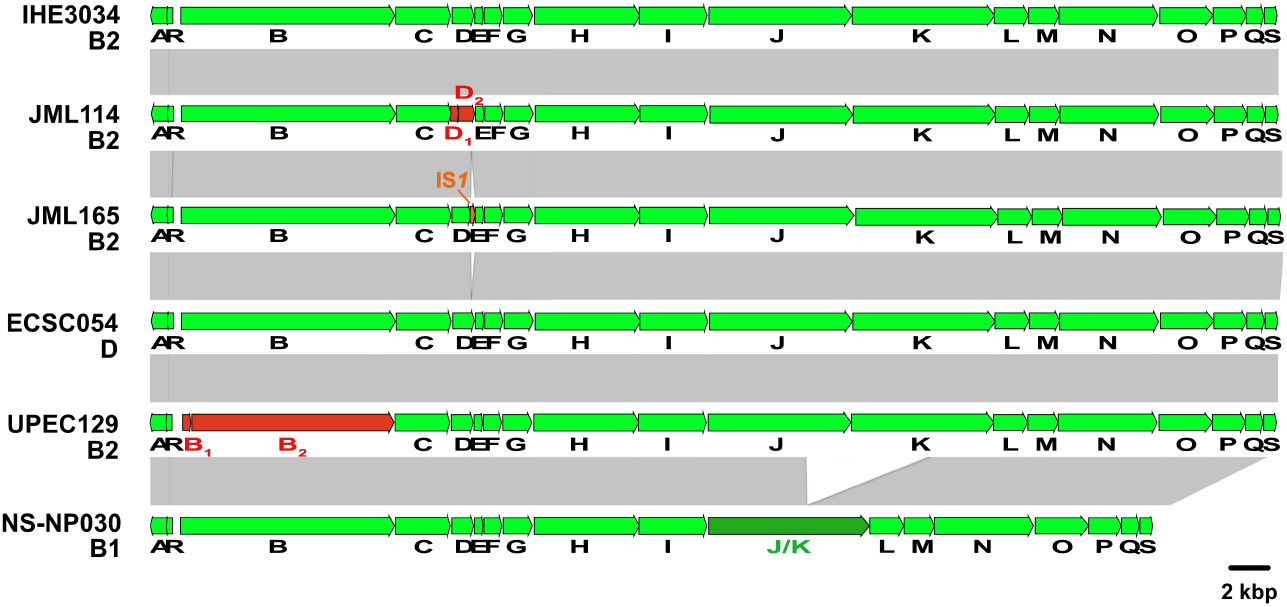
Comparison of the *pks* island sequence from the *E. coli* reference strain IHE3034 with that of a selection of *pks-*positive but non-genotoxic *E. coli* isolates. Nucleotide sequence similarity (>99%) between different DNA regions is indicated by grey areas between the corresponding genetic maps. Fusion of two adjacent ORFs resulting from the deletion of a sequence overlapping the two ORFs is indicated in dark green. Adjacent ORF sequences resulting from the segregation of an original ORF following an insertion or deletion event are indicated in red. The IS*1* located in the *pks* island of strain JML165 is represented in orange.

Twelve ICL-negative strains carried a 5,651-bp deletion resulting in a *clbJK* fusion gene, as shown for strain NS-NP030 in Fig. 7. This deletion presumably resulted from recombination between two copies of a 1,480-bp homologous sequence located in *clbJ* and *clbK*. A PCR analysis of the corresponding region in the 109 *pks*-positive strains confirmed the presence of this deletion in the 12 strains, whereas the other 97 *pks*-positive strains contained full-length *clbJ* and *clbK* genes (Fig. 5). The 12 strains carrying the *clbJK* fusion were detected sporadically in the core genome phylogenetic tree (Fig. 1) and *pks* phylogenetic tree (Fig. 5), suggesting that occurrence of the deletion between *clbJ* and *clbK* arose from accidental recombination events. The predicted 2,440-amino acid hybrid ClbJK protein encoded by the *clbJK* fusion gene lacks the PKS module of ClbK necessary for the formation of stable cross-links (19). In agreement with this, the strains carrying this fusion were devoid of interstrand crosslinking activity and did not trigger megalocytosis nor histone H2AX phosphorylation in infected eukaryotic cells (Table 3). It was reported however that rat *E. coli* isolates carrying a *clbJK* fusion gene caused DNA damage or displayed cytotoxicity to HeLa cells (39, 40). This discrepancy with our results could be due to the use of distinct experimental conditions. The possibility that these rat isolates might produce additional genotoxins that would mask any colibactin deficiency caused by the *clbJK* fusion cannot be excluded either. Caution should also be observed during the assembly of sequencing reads as errors including deletions may be caused by the presence of tandem repeats.

For two other non-genotoxic isolates (CM1 and ECSC054), no mutation disrupting the *pks* genes were identified (Fig. 7; data not shown), suggesting that mutations located outside the *pks* island could negatively impact its expression. To test this hypothesis, we used plasmids pASK-clbR and pBAD-clbR, both overexpressing the *pks* regulator ClbR, and introduced either of them into the strain CM1 which was susceptible to antibiotic, in contrast to ECSC054. In the resulting CM1 transformants, colibactin activity was restored as seen by the formation of DNA ICLs (data not shown). Thus, in this strain, the lack of genotoxic activity likely resulted from a negative regulation of the *pks* island through an unknown mechanism.

Functionality of the *pks* island was also examined through analysis of the lipid metabolite profiles of selected genotoxic (n=3) and non-genotoxic (n=8) *pks*-positive strains, and in particular for production of C14-Asn which was used as an indicator of the activity of the *pks* biosynthesis machinery. This lipopeptide is synthesized during the initial step of the biosynthesis process involving ClbN and ClbB, prior to elongation and final cleavage through the involvement of ClbC-H-I-J-K and ClbP, respectively (17). The production of C14-Asn was detected in all of the ICL-positive strains examined, SI-NP020, JML285 and KS-NP019 (*ca*. 650-1200 pg/10e8 CFU) but not in the ICL-negative strain UPEC129 mutated in *clbB* (Fig. S5, Table 3). Interestingly, C14-Asn was detected (*ca*. 400-600 pg/10e8 CFU) in three ICL-negative strains carrying a *clbJK* fusion gene (SI-NP032, JML296 and NS-NP030) and in two ICL-negative strains carrying a mutated *clbD* gene (JML114 and JML165). The two non-genotoxic strains carrying intact *clb* genes (ECSC054 and CM1) produced either a very low level or no detectable C14-Asn, respectively (Fig. S5, Table 3). For the strain CM1 transformed with either plasmid pASK-clbR or pBAD-clbR (see above), overexpression of ClbR restored the production of C14-Asn (Fig. S5). These results suggest that even when the *pks* island does not allow production of active colibactin, enzymes from the *pks* pathway still produce metabolites with potential biological activities.

## DISCUSSION

The acquisition of the *pks* island in the population of *E. coli* appears to have involved two distinct mechanisms differing by the presence or absence of a phage-type integrase. The integrase-mediated *pks* insertion pathway occurred mainly in B2 strains and resulted in *pks* insertion into either of three *asn* tRNA genes (i.e. *asnU, asnV* or *asnW*). This potential for integration into several DNA targets is consistent with the observed conservation and genetic integrity of the *pks* integrative module, i.e. the integrase gene and the two direct repeats flanking the island. The flexibility of *pks* insertion is reminiscent of what has been described for the HPI of *Yersinia pseudotuberculosis* which is also able to insert into either of the three *Y. pseudotuberculosis asn* tRNA genes (41), in contrast to the immobile truncated form of the HPI in *E. coli* whose right direct repeat is deleted and whose location is fixed at the *asnT* tRNA gene (42). A divergent integrase sequence was found for the *pks* island inserted into the *asnU* tRNA gene compared to those inserted into the *asnV* or *asnW* tRNA gene. As the three *asn* tRNA sequences are 100% identical, the use of either of them as attachment site by slightly different integrases likely reflects distinct histories of *pks* acquisition. After *pks* chromosomal integration, the endogenous *pks* integrase promoter is replaced by the promoter of the upstream *asn* tRNA gene (Fig. S6), a configuration similar to that found for the HPI integrase promoter (43). Whether the site of integration influences the expression of the *pks* integrase and hence *pks* stability at the distinct *asn* tRNA *loci* is not known. In contrast to the integrase-mediated pathway, the *pks* chromosomal integration process in the B1 and A *E. coli* strains remains unclear as no site-specific recombinase-encoding gene was found near *pks* and chromosomal insertion occurred into a non-tRNA *locus*. In these strains, *pks* integration could have involved the participation of IS elements such as the IS*66* whose truncated or intact copies were found to flank the *pks* island.

The cophylogeny analysis between the core genome- and *pks*-based phylogenetic trees shed further lights on the *pks* acquisition scenarios. In the case of the typical *pks*-positive B2 strains belonging to lineages from major CCs (i.e. CC12, CC14, CC73 and CC95), the congruence observed between both trees suggested that the *pks* island was horizontally acquired by the MRCA of these lineages, or by the MRCA of each of these, and then stably maintained in their descendants through vertical transmission. The fact that strains of certain CC95 subgroups lack the *pks* island likely suggests that *pks* was lost during the evolution of these sublineages. Such a loss might be closely linked to the change in the relative fitness of CC95 subgroups underlying the variations observed in their spatial and temporal distribution in several continents (35). In the case of B1 or A strains, *pks* acquisition and dissemination likely occurred through sporadic lateral transfer events, as *pks*-positive B1 or A strains were scarce and not genetically related. The horizontal transferability of the *pks* island has previously been demonstrated using an *in vitro* approach where *pks* could be transferred together with the HPI *via* F’ plasmid-mediated conjugation from a donor to a recipient *E. coli* strain (44). We propose that *pks* acquisition by the single *pks*-positive D strain ECSC054 was mediated by HGT, presumably from a B2 donor strain given the *pks* sequence relatedness observed between the D and B2 strains. HGT was also likely involved in the exchange of *pks* island between the three atypical B2 ST73 or ST95 strains and a (yet unknown) phylogenetically distant donor strain, since their *pks* sequences did not cluster with those from other B2 ST73 or ST95 strains. This hypothesis was further supported by the identification, in these three isolates, of an ICE-like element inserted in their *pks* island. Similar ICE-like elements have previously been identified in three B1 *E. coli* isolates and other members of the *Enterobacteriacae* such as *C. koseri, E. aerogenes* and *K. pneumoniae* (6). They could therefore play a role in *pks* dissemination in enterobacteria, as proposed for the self-transmissible ICE linked to the HPI identified in the *E. coli* strain ECOR31 (38). Due to its lack of a complete DNA mobilization region (region II), we assume that the ICE-linked *pks* island is not self-transferable anymore. It might nevertheless correspond to a remnant of an ancient, complete and self-transmissible ICE-linked *pks* island that could have behaved as a large complex ICE and spread in enterobacteria before undergoing partial or entire deletion of the ICE region. To date, no bacterial strain carrying a complete ICE linked to *pks* has been identified and the origin of the *pks* island remains therefore elusive.

The ecological niche and/or genetic background of the bacterial strains probably had an impact on the acquisition and stable maintenance of *pks*. The high concentration of *pks*-positive strains in some CCs of the B2 group such as CC73 and CC95 suggests that *pks* might have contributed to their ecological and evolutionary success. CC73 and CC95 exhibit a similar phylogenetic history and are major ExPEC lineages, especially prior to the year 2000 where they were the most commonly detected (1, 45). They are persistent intestinal colonizers and successful extra-intestinal pathogens with the particularity of exhibiting lower multidrug resistance levels compared to other ExPEC lineages. By contrast, as our collection contained *E. coli* B2 strains from STs other than ST73 and ST95, it was interesting to note that *pks* was absent from STs corresponding to separate B2 lineages in the *E. coli* phylogenetic tree, including the ST131 clonal complex which is associated with multidrug resistance and is now the most predominantly isolated ExPEC lineage worldwide (45). Consistent with the hypothesis of a *pks* acquisition by the MRCA(s) of CC73 and CC95 mentioned above, this finding suggests that such acquisition likely occurred after they diverged from the MRCA of CC131, i.e. before going through distinct evolutionary trajectories. The inversion of the upstream *asnW-asnU-asnV* tRNA-containing region which likely accompanied *pks* insertion into the *asnW* tRNA gene in the B2 group might have contributed to *pks* stabilization at this locus. Since the various *pks*-positive and -negative B2 lineages occupy the same ecological niche (i.e. primarily the intestinal tract of humans and animals), horizontal transfer of *pks* between them could have been expected, at least to some extent, which however was not revealed here. Several hypotheses can be proposed to explain this. First, some barriers to HGT might exist between members of distinct CCs, such as restriction-modification systems (46). Second, *pks* might have been transferred to recipient strains without providing adaptive value, thus resulting in its rapid loss. Third, as a crosstalk between virulence determinants and the chromosome backbone is required for the emergence of virulent clones (1), a specific chromosomal phylogenetic background might be required for appropriate *pks* expression and production of an adaptive value, thereby constituting a prerequisite to the stable maintenance of the island.

The structure of the *pks* island is very well conserved among the *E. coli* population, with more than 99 % identity, suggesting that its integrity remains under strong structural and functional evolutionary constraints. We can speculate that, transcription and translation of the 19 *pks* genes of this 54 kb-long genomic island would be too high for the bacterial strains if the *pks* island did not bring a selective advantage to them. This is reinforced by the fact that only 18 out of 109 *pks*-positive strains lacked genotoxic activity. The importance of *pks* biological role is highlighted by the numerous activities associated with this genetic island, including genotoxicity, anti-inflammatory activity, antibiotic and analgesic effects. Given its interplay with siderophores (enterobactin, salmochelin and yersiniabactin) and siderophore-microcins (MccM and MccH47) (10, 22, 47), the *pks* island contributes to bacterial competition through the acquisition of iron or the production of inhibitory compounds, respectively. Protection of bacterial cells from genomic degradation through the production of ClbS could also be advantageous to *pks*-carrrying strains, as this multifunctional protein not only directly inactivates colibactin but also protects bacterial DNA from nucleolytic degradation by nucleases (48). We also observed that non-genotoxic *E. coli* strains carrying an altered *pks* island still produced the prodrug motif C14-Asn synthesized at the early stage of the biosynthesis process, suggesting that yet-to-be-discovered bioactive compounds are produced by these strains. Given the high conservation observed for the *pks* island in *E. coli*, we can thus speculate that colibactin is a very important genotoxin but that *pks*-derived synthesis of other secondary metabolites could also be an advantage for *E. coli*.

Although our collection is characterized by a large diversity of *E. coli* strains from various phylogenetic groups and STs, one limitation of this study is that only strains from Japan were included, which may not be representative of *pks* distribution in a global collection of *E. coli* isolates from worldwide sources.

In conclusion, the various genetic configurations of the *pks* island and its distribution in the *E. coli* phylogenetic tree imply the existence of various scenarios for the introduction and spread of *pks* into the *E. coli* population. The presence of a functional *pks* island was demonstrated for the majority of the *pks-*positive strains, suggesting that the *pks* island is under selective pressure for the adaptation of *E. coli* to various ecological niches, through the production of colibactin or other secondary metabolites.

## AUTHOR STATEMENTS

### Author contributions

Conceptualisation: FA, TH, PB, YO, EO. Methodology: FA, TH, PB, YO, EO. Validation: FA, TH, PB, YO, EO. Formal Analysis: FA, AP, CC, YA, TH, PB, YO, EO. Investigation: FA, AP, YA, CC, NBG, CM, JPN, PB, YO, EO. Resources: TH, YO, EO. Data Curation: FA, AP, YA, TH, PB, YO, EO. Writing – Original Draft: FA, TH, YO, EO. Writing - Review and Editing: FA, AP, CC, HB, JPN, TH, PB, YO, EO. Visualization: FA, YA, CC, JPN, PB, YO, EO. Supervision: FA, HB, TH, YO, EO. Project Administration: FA, EO. Funding Acquisition: TH, YO, EO.

## Acknowledgments

We thank Claire Hoede and Sarah Maman (SIGENAE group) and the GENOTOUL bioinformatics platform for providing computational resources. We also thank Pauline Le Faouder and the METATOUL lipidomic platform for their support in the analysis of the lipid metabolite profiles.

