## Supplementary material for "Insights into the acquisition of the *pks* island and production of colibactin in the *Escherichia coli* population": Fig. S

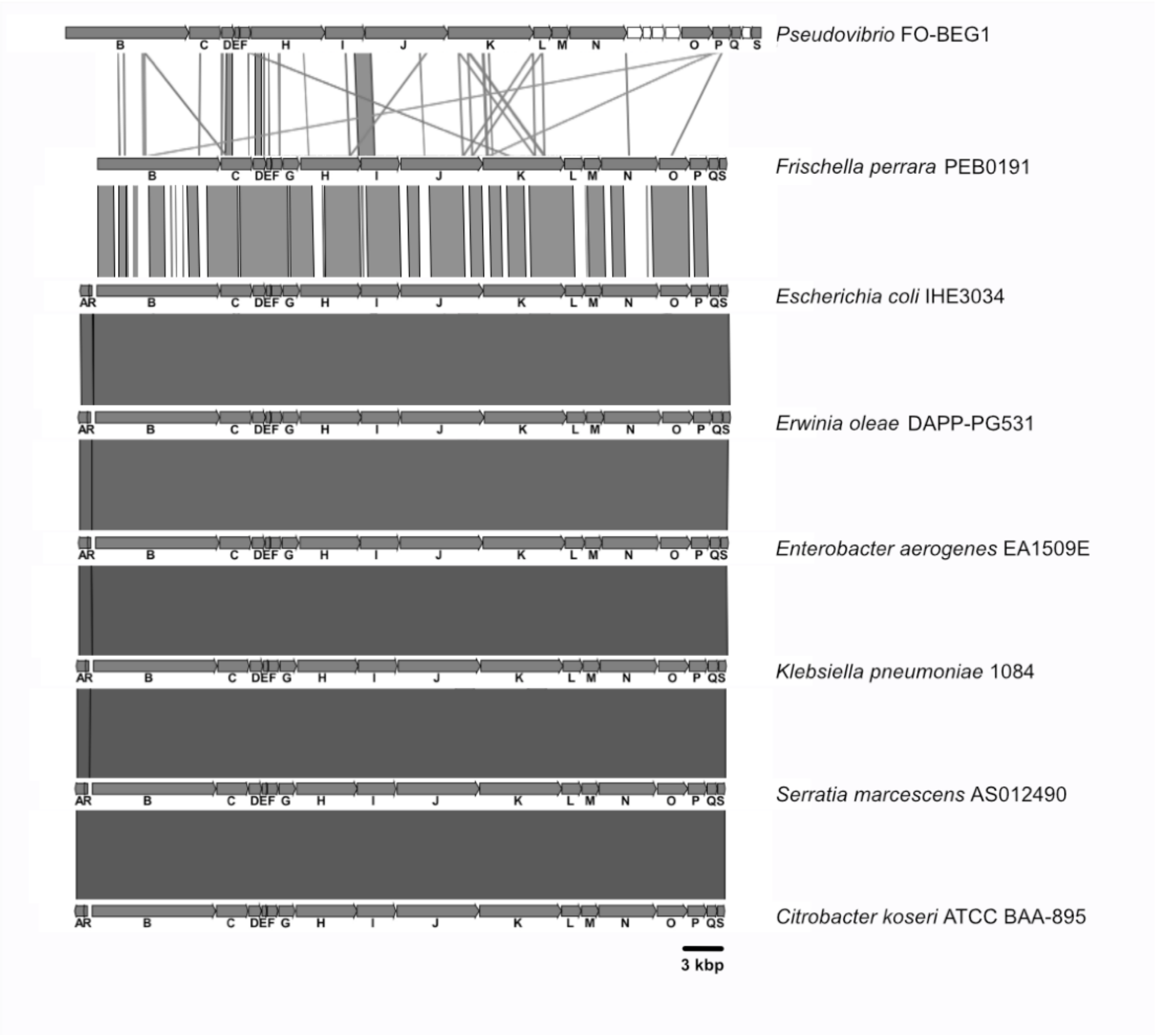


**Figure S1.** Comparison of the *pks* island of reference *E. coli* strain IHE3034 with that of non-*E. coli* strains. The non-*E. coli* strains included 5 members of the *Enterobacteriaceae* family (*C. koseri* ATCC BAA-895, *E. aerogenes* EA1509E, *K. pneumoniae* 1084, *S. marcescens* AS012490 and *E. oleae* DAPP-PG531), the honey bee strain *F. perrara* PEB0191 (family of the *Orbaceae*) and the marine alphaproteobacterial strain *Pseudovibrio* FO-BEG1. Nucleotide sequence similarity (>99%) between different DNA regions is indicated by gray areas between the corresponding genetic maps.

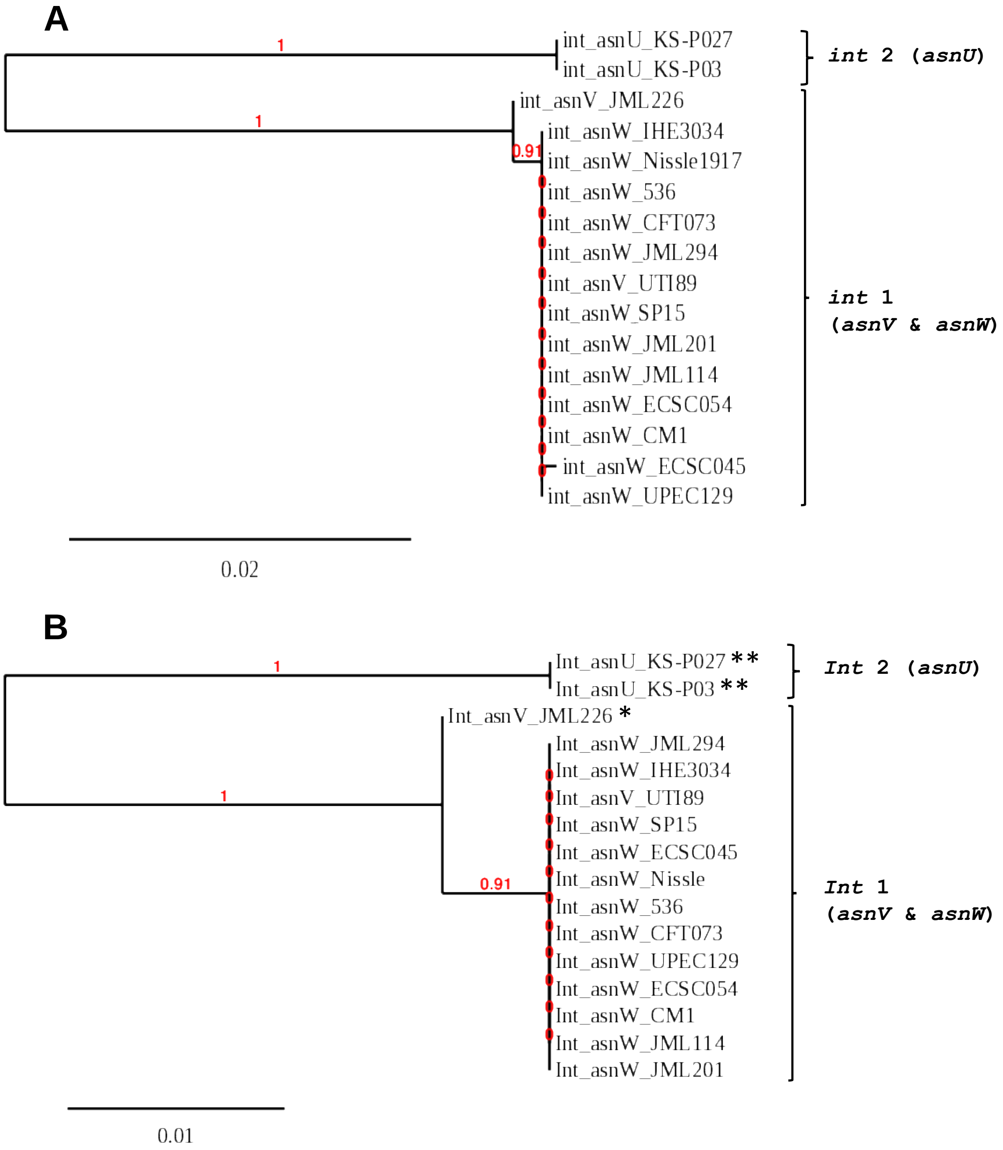


**Figure S2.** Phylogenetic tree of the nucleotide (A) and amino acid (B) sequences of *pks* integrases encoded by 16 *pks*-positive *E. coli* strains, carrying an integrase gene at either the *asnU*, *asnV* or *asnW* tRNA gene. All the strains belong to the B2 phylogroup, except ECSC054 (phylogroup A). The B2 strains were selected to include both isolates (*i.e.* KS-P03 and KS-P027) carrying *pks* at the *asnU* locus, both isolates (*i.e.* JML226 and UTI89) carrying *pks* at the *asnV* locus and additional B2 isolates of ST73 (i.e. CFT073, JML114 and Nissle), ST95 (i.e. CM1, ECSC045, JML201, JML294, SP15 and UPEC129) and 127 (536). The type (U, V or W) of the *asn* tRNA gene located upstream the integrase-encoding gene is indicated. The two clusters of integrase sequences and the corresponding integrase families (Int1 and Int2) are shown on the right. Branch support values are indicated in red. *, two amino acid substitutions; **, 23 amino acid substitutions, compared to the integrase from reference strain IHE3034.

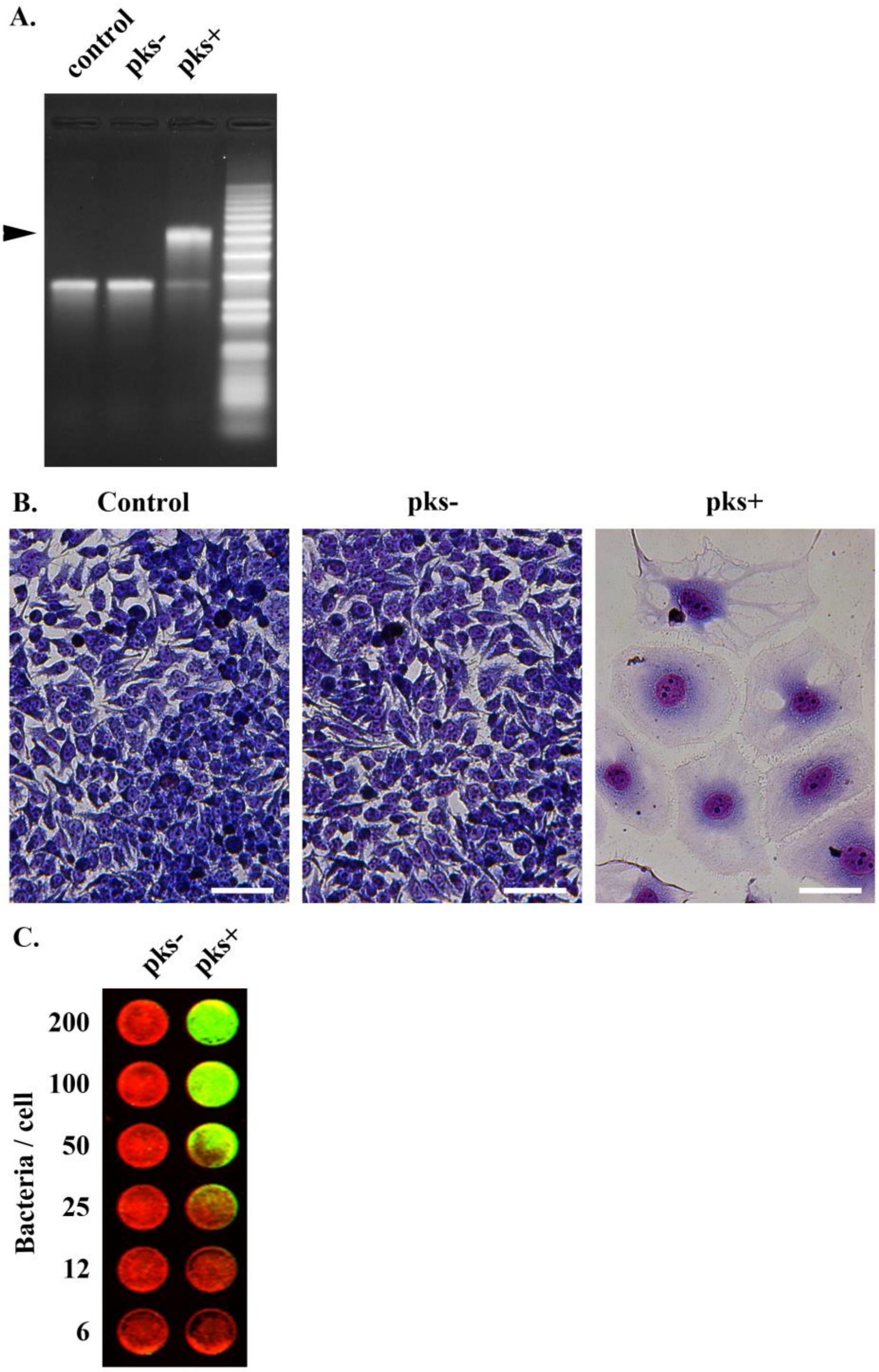


**Figure S3.** Functionality testing of the *pks* island. **(A)** *In vitro* DNA interstrand cross-linking assay. Linearized double-stranded plasmid DNA was exposed to DH10B pBACpks (pks+) or vector (pks-) for 40 min at 37°C and then analyzed by denaturing electrophoresis. The arrowhead indicates the interstrand crosslinked DNA with a two-fold lower electrophoretic mobility. **(B)** Megalocytosis (cell enlargement) assay. HeLa cells were infected 4 h with either *pks*-negative or *pks*-positive *E. coli* strains at a multiplicity of infection of 400 bacteria per cell. Cells were then washed and incubated with gentamicin for 48-72 hours before staining with methylene blue. Bars, 50 µm. **(C)** *In cellulo* DNA damage assay. HeLa cells were transiently infected with either *pks*-negative or *pks*-positive *E. coli* at various multiplicity of infection, followed by washes and incubation with gentamicin. Cellular DNA damage was revealed by staining of p-H2AX (green) relative to nuclear DNA (red).

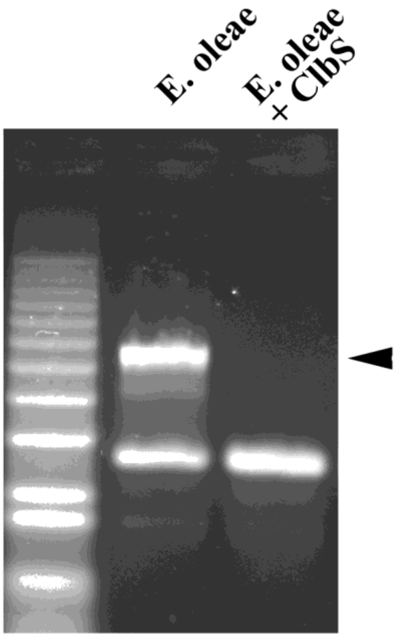


**Figure S4.** Functionality testing of the *pks* island of *E. oleae* strain DAPP-531. Linearized double-stranded plasmid DNA was exposed to *E. oleae* strain DAPP-531 for 40 min at 37°C in the absence (*E. oleae*) or presence of 400nM purified His-tagged ClbS protein (*E. oleae* + ClbS), and then analyzed by denaturing gel electrophoresis. The arrowhead indicates the interstrand crosslinked DNA with lower electrophoretic mobility.

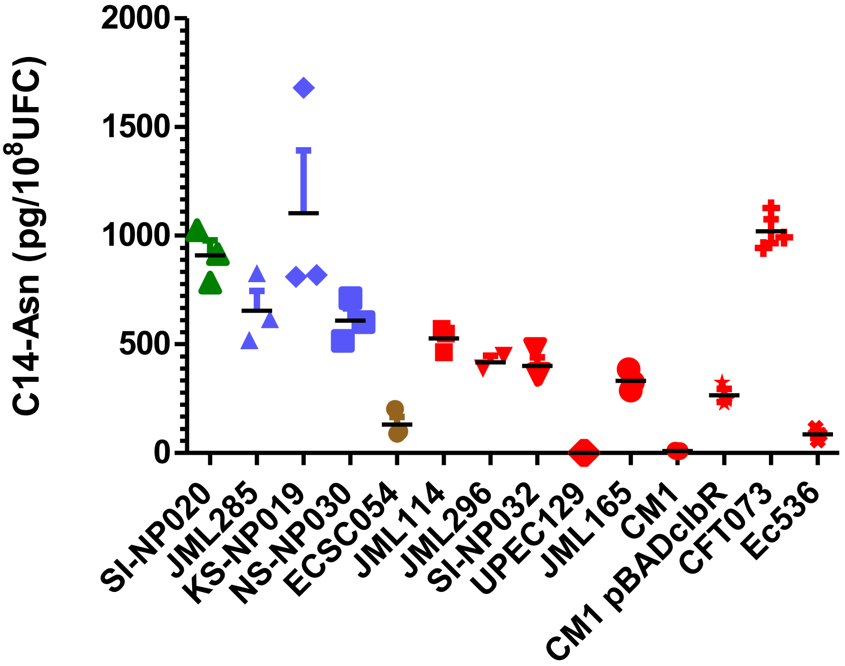


**Figure S5.** Quantification of C14-Asparagine from *pks*-positive *E. coli* strains. C14-Asn was quantified for *E. coli* strains belonging to phylogroups A (green), B1 (blue), D (brown) and B2 (red), including control B2 strains CFT073 and Ec536. Data are represented as mean +/- SEM of 3 independent bacterial cultures per group.

**A.**

-35

ACGACAAAAT GCAGATTATT TCAGCAAACG ATTTCAAATT TAAAAAACAG GCTTTGACAT

-10

TGTGGGTGGG CATCGCTAAT ATTCGCCTCG TTCTCACGAT *TCCTCTGTAG TTCAGTCGGT*

*AGAACGGCGG ACTGTTAATC CGTATGTCAC TGGTTCGAGT CCAGTCAGAG GAGCCA*ATTT

CCTGCTTTCA TGCATCCTTG CGAATCCTTA TGTTATTTTT ATTCAACAGG TTAGCGTGAA

AACTCTTCCC GGTGCATTTT GATTTTACCC TCTGCATCGG GAAAAATTGG TAGTCAAATC

TGGGGTCAGG TTAGTTCGAT AATGGAGTGA CCCCCAT**ATG** TCCCTTAACG ACGCAAAAAT

CCGTAGTCTC AAGCCCACTG ATAAACCCTT TAAAGTCTCC GATTCCCACG GTCTGTATCT

**B.**

-35

TTTATGTAGC CAGCTCCTAT TGGTGGTCAT TCTGGTGGTC TTGACAGGAA GATAACTCTG

-10 +1

GTTTAGCTTA CTTATTAGCC ACTTACTGGC AAAGGCGATC CCAGTCAGAG GAGCCAATTT

CCTGCTTTCA TGCATCCTTG CGAATCCTTA TGTTATTTTT ATTCAACAGG TTAGCGTGAA

AACTCTTCCC GGTGCATTTT GATTTTACCC TCTGCATCGG GAAAAATTGG TAGTCAAATC

TGGGGTCAGG TTAGTTCGAT AATGGAGTGA CCCCCAT**ATG** TCCCTTAACG ACGCAAAAAT

CCGTAGTCTC AAGCCCACTG ATAAACCCTT TAAAGTCTCC GATTCCCACG GTCTGTATCT

**Figure S6.** Analysis of the promoter region of the *pks* integrase gene. **(A)** Nucleotide sequence of the promoter region of the *pks* integrase gene from reference strain IHE3034. The *asnW* tRNA gene is indicated in italics and its -10 and -35 promoter hexamers are underlined. The 17-bp core sequence where site-specific recombination occurs is boxed. The start codon (ATG) of the integrase gene is in bold letters. **(B)** Nucleotide sequence of the promoter region of the *pks* integrase gene reconstituted after theoretical chromosomal excision of the island. The putative -10 and -35 promoter hexamers and transcriptional start (+1) of the integrase gene are underlined and the start codon (ATG) is in bold letters. The 17-bp core sequence is boxed.
